# Multiple ancestral and a plethora of recent gene duplications during the evolution of the green sensitive opsin genes (*RH2*) in teleost fishes

**DOI:** 10.1101/2021.05.11.443711

**Authors:** Zuzana Musilova, Fabio Cortesi

**Affiliations:** Department of Zoology, Faculty of Science, Charles University, Vinicna 7, 12844 Prague, Czech Republic; Queensland Brain Institute, The University of Queensland, Brisbane, QLD 4072, Australia

## Abstract

Vertebrates have four visual cone opsin classes that, together with a light-sensitive chromophore, provide sensitivity from the ultraviolet to the red wavelengths of light. The rhodopsin-like 2 (RH2) opsin is sensitive to the centre blue-green part of the spectrum, which is the most prevalent light underwater. While various vertebrate groups such as mammals and sharks have lost the *RH2* gene, in teleost fishes this opsin has continued to proliferate. By investigating the genomes of 115 teleost species, we find that *RH2* shows an extremely dynamic evolutionary history with repeated gene duplications, gene losses and gene conversion affecting entire orders, families and species. At least four ancestral duplications provided the substrate for today’s *RH2* diversity with duplications occurring in the common ancestors of Clupeocephala, Neoteleostei, and Acanthopterygii. Following these events, *RH2* has continued to duplicate both in tandem and during lineage specific genome duplications. However, it has also been lost many times over so that in the genomes of extant teleosts, we find between zero to eight *RH2* copies. Using retinal transcriptomes in a phylogenetic representative dataset of 30 species, we show that *RH2* is expressed as the dominant green-sensitive opsin in almost all fish lineages. The exceptions are the Osteoglossomorpha (bony tongues and mooneyes) and several characin species that have lost *RH2*, and tarpons, other characins and gobies which do not or only lowly express the gene. These fishes instead express a green-shifted long-wavelength-sensitive *LWS* opsin. Our study highlights the strength of using modern genomic tools within a comparative framework to elucidate the detailed evolutionary history of gene families.

## Introduction

Vertebrates use vision to find food and mates, avoid predators and to navigate their surroundings. At the centre of their visual systems lie the visual pigments which consist of an opsin receptor protein that is covalently bound to a vitamin A derived, light sensitive chromophore (Hunt et al., 2014; Wald, 1968). Five opsin types were already present in the vertebrate ancestor which can be distinguished by their photoreceptor specificity, their phylogeny and their maximum light sensitivity (λ_max_) (Lamb, 2013). The rod or rhodopsin (RH1) is sensitive to the blue-green light spectrum ~ 450 – 540 nm λ_max_ and is expressed in the highly sensitive rod photoreceptors, which are active during dim light. Four cone-specific opsins are active during bright light conditions and confer spectral sensitivity from the ultraviolet (UV) to the red spectrum. These are the short-wavelength-sensitive 1 (SWS1, 345 – 440 nm λ_max_), SWS2 (395 – 490 nm λ_max_), rhodopsin-like 2 (RH2, 450 – 540 nm λ_max_), and the long-wavelength-sensitive (LWS, 490 – 575 nm λ_max_) (reviewed in Carleton et al., 2020; Yokoyama, 2008). As opposed to terrestrial vertebrates where the ancestral five opsins have either been maintained (e.g., birds) or some types might have been lost (e.g., mammals) (Hunt et al., 2014), in teleost fishes opsin genes have continued to proliferate at an astonishing rate (reviewed in Musilova et al., 2021). Amongst the cone opsins, *RH2* appears to have the highest number of gene duplicates in teleosts, with eight copies found in squirrel and soldierfishes (Holocentridae) (Musilova et al., 2019).

RH2-based visual pigments are sensitive to the middle of the light spectrum (blue to green) (Yokoyama and Jia, 2020), which is the most commonly available light underwater, especially with increasing depth (Jerlov, 1976; Munz and McFarland, 1977). As light is quickly scattered and absorbed in aquatic environments, deeper bodies of water lack shorter and longer wavelengths of light with fishes in those habitats often lacking the UV sensitive *SWS1* and red sensitive *LWS* genes but instead they have an increased number of *RH2* duplicates (Musilova et al., 2019). Similarly, species that are active during the night or during twilight hours, where blue to green light dominates, show increased reliance on *RH2* genes. As for the deeper living species, crepuscular and nocturnal fishes have often lost or do not express the genes sensitive to the edges of the visible light spectrum such as found in coral reef squirrel and soldierfishes (Busserolles et al., 2021) and cardinalfishes (Apagonidae) (Luehrmann et al., 2019). *RH2* and *RH1* share a common ancestry (reviewed in Musilova et al., 2021), and in the deep-sea pearlsides (*Maurolicus* spp.) *RH2* is expressed in rod-like cone cells (de Busserolles et al., 2017). Pearlsides are active during dusk and dawn when they come to the surface to feed while for the remainder of the time, they sink to a depth of around 200 m to rest (Giske et al., 1990).

It is during these twilight hours, when the light environment is blue shifted and at an intensity that is neither ideal for cone nor rod cells, that the transmuted rod-like cones seem to function at their best (de Busserolles et al., 2017). In the extreme case of deep-sea fishes, *RH2* is often the only remaining cone opsin gene in their genomes, where it is mostly expressed in the larval stages that start their lives in the shallow and well-lit epipelagic ocean (Lupše et al., 2021).

Interestingly, in the Osteoglossomorpha (bony tongues and mooneyes) and several characin species, *RH2* has been lost over evolutionary time and green sensitivity has been acquired by a modified, shorter-wavelength sensitive copy of the *LWS* opsin gene (Escobar-Camacho et al., 2020; Liu et al., 2018). This is similar to what occurred in primates including humans, whereby *RH2* was lost in the mammalian ancestor and green vision was recovered by means of a second, mid-wavelength sensitive (MWS) *LWS* duplicate (Carvalho et al., 2017; Yokoyama and Yokoyama, 1990). In gobies and most characins, *RH2* is still present, but nonetheless green sensitivity has been taken over by shorter shifted *LWS* duplicates (Cortesi et al., 2021; Escobar-Camacho et al., 2020). This shows that the spectral sensitivity range and function for specific types of opsins might not be as conserved as was previously assumed, calling for a more thorough investigation of this diverse gene family.

To understand the function of *RH2* more thoroughly and gain a comprehensive overview of the dynamics underlying the *RH2* evolution in fishes, we set out and mined publicly available genomes from a phylogenetic representative dataset of 115 teleost species. In doing so we uncover a rich evolutionary history that is characterised by multiple ancestral duplications and a plethora of more recent duplications, gene losses and gene conversion. Combined with retinal transcriptomes from 30 species, we show that *RH2* is expressed in the majority of fish species, but we also uncover a number of cases where its function has been taken over by other opsin genes, namely *LWS*.

## Results and Discussion

Mining the genomes of 115 teleost species (and three non-teleost outgroups) and reconstructing the most extensive *RH2* phylogeny to date, we find that ray-finned fishes possess a median number of three *RH2* copies per species, thus making it the most numerous opsin gene in teleost fishes (Figs. 1 and 2, and Fig. S1). We also confirm that the blackbar soldierfish, *Myripristis jacobus*, with eight *RH2*s, has the highest number of paralogs in its genome (also see Musilova et al., 2019). Moreover, several species were found to have up to seven *RH2* duplicates including the milkfish (*Chanos chanos*) and the glacier lanternfish (*Benthosema glaciale*). However, other than the previously reported Osteoglossomorpha (Liu et al., 2018) and several characin species (Escobar-Camacho et al., 2020), we did not find any species that has lost *RH2* altogether (Fig. 1). Instead, based on retinal transcriptomes in 30 species, we discovered several species and entire lineages where *RH2* is not or only minimally expressed, at least in the adult stages (Fig. 3).

**Figure 1.**
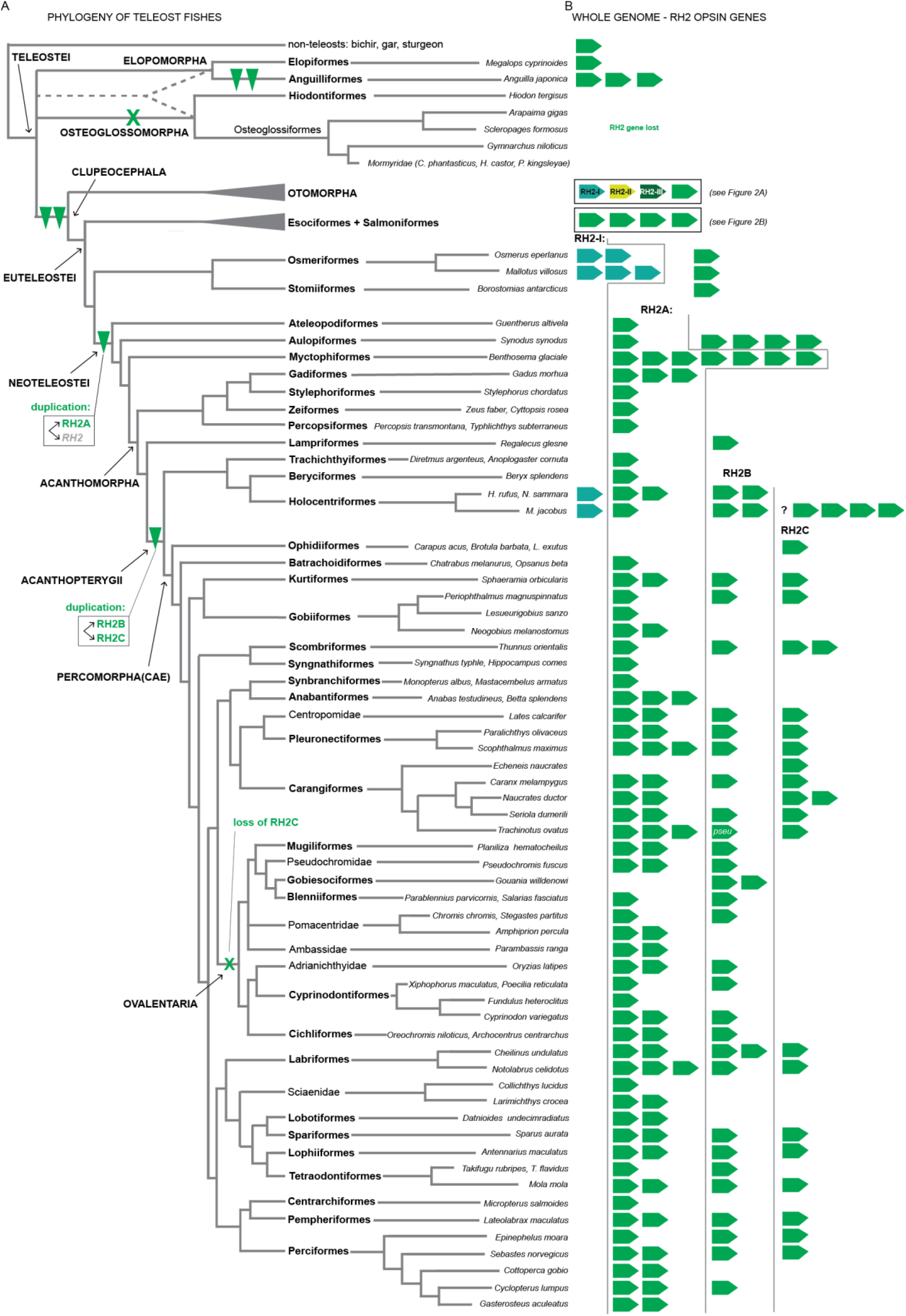
Evolution and diversity of the *RH2* opsin genes in teleost fishes. A) Schematic phylogeny (after Betancur-R et al., 2017) of the teleost genomes that were analysed in this study. At least two *RH2* gene duplications occurred in the ancestor of Clupeocephala giving rise to the *RH2-I*, *RH2-II*, *RH2-III*, and *RH2* copies. Further duplications occurred later in the ancestor of Neoteleostei (giving rise to *RH2A*) and Acanthopterygii (giving rise to *RH2B* and *RH2C*). Because there is an uncertainty with the assignment of the holocentrid *RH2* genes, the alternative scenario is that this duplication happened in the ancestor of Percomorpha. The *RH2* opsin gene type was lost in Osteoglossomorpha and a few characins (see Escobar-Camacho et al., 2020) for details on characins) the *RH2C* copy was lost in Ovalentaria. B) Number and types of the *RH2* opsins found in the whole genomes of 115 ray-finned fishes. See Fig. 2 for details on the Otomorpha, Esociformes, and Salmoniformes *RH2* evolution. RH2, rhodopsin-like 2 opsin gene; pseu, pseudogene.

**Figure 2.**
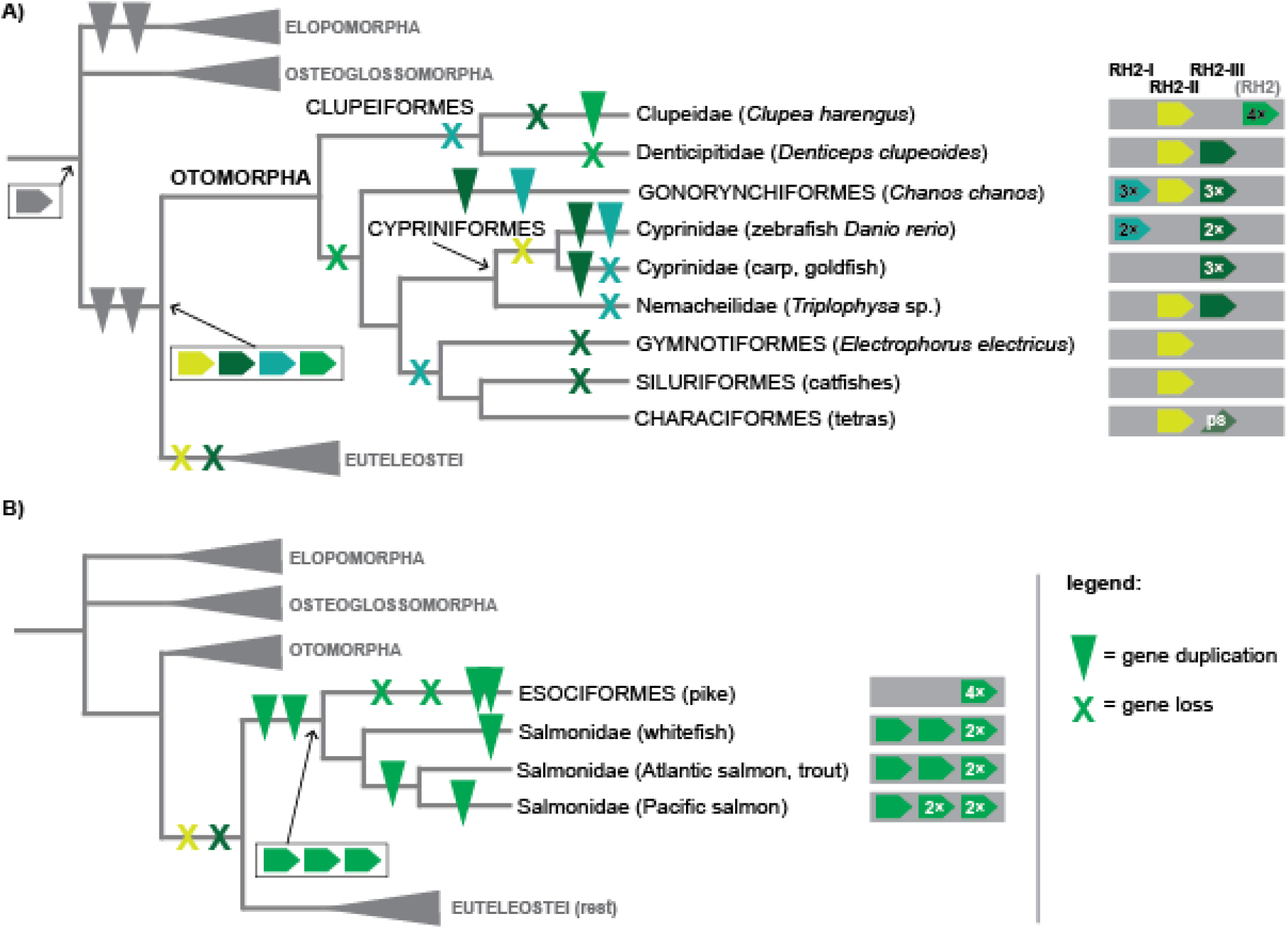
Evolution of the *RH2* opsin genes in Otomorpha, Esociformes, and Salmoniformes. A) In Otomorpha the *RH2* diversity is based on all four ancestral copies. The *RH2-II* and *RH2-III* copies persisted only in Otomorpha, *RH2-I* is also preserved in smelts (Osmeriformes) and soldierfishes (Holocentriformes) (see Fig. 1). The *RH2-I* and *RH2-III* copies are located in the conventional *RH2* cluster flanked by the *synpr* and *slc6A13* genes, similar to the euteleost *RH2s* (data not shown). The *RH2-II* copy is located on the separate cluster next to the *mut5-HS* gene (data not shown). Note, for example, that the only *RH2* gene in catfishes and characins (*RH2-II*) is a different paralog than the copy used in cyprinids (*RH2-III*). B) In pike and salmon, *RH2* duplicated from the fourth ancestral copy, likely in their common ancestor and these paralogs are not shared with other teleost lineages. RH2, rhodopsin-like 2 opsin gene.

**Figure 3.**
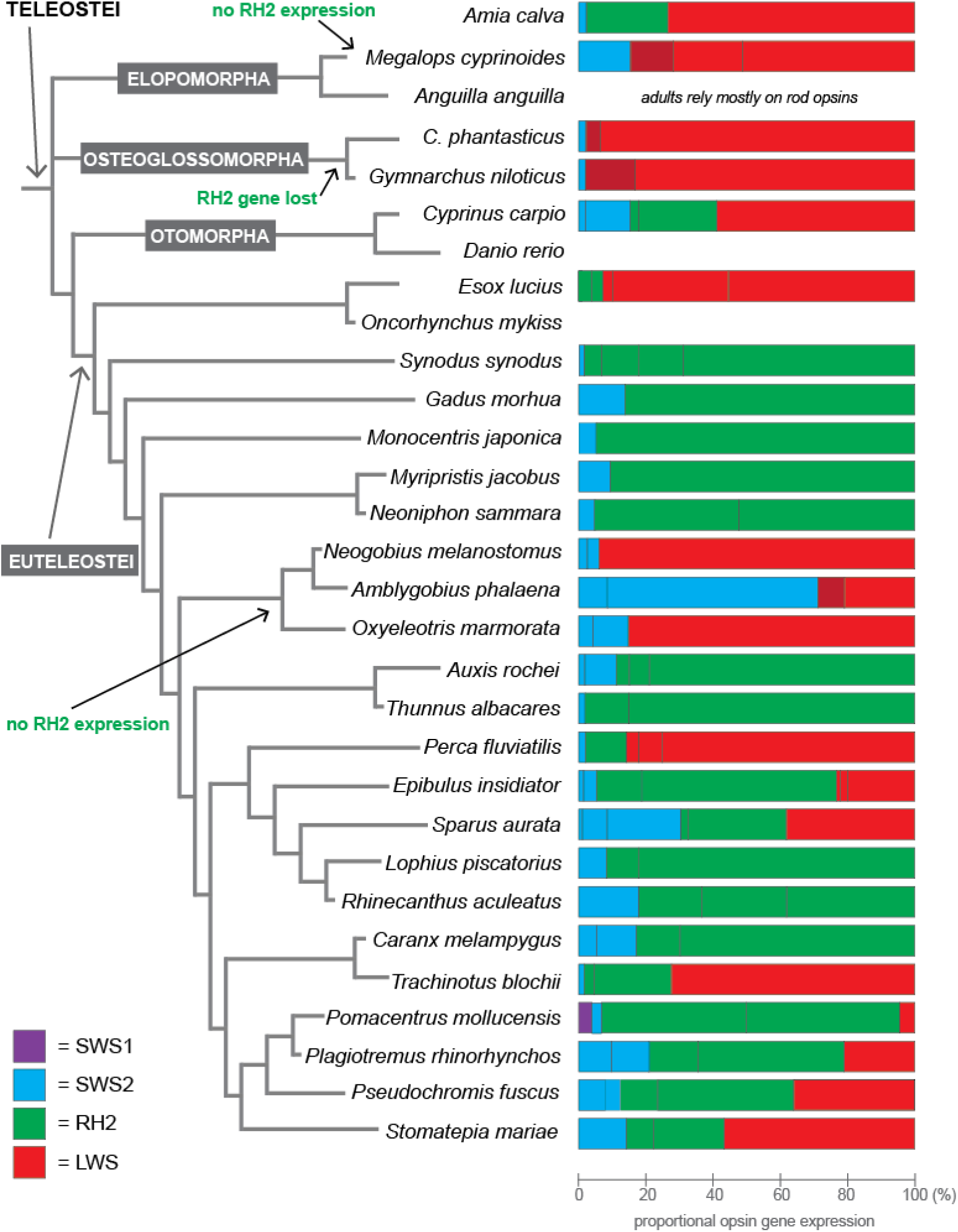
Cone opsin gene expression in teleost fishes. The *RH2* opsin gene is expressed in most of the species highlighting its relevance for vision in different aquatic habitats (from deep sea to shallow streams). *RH2* has been lost in Osteoglossmorpha and none or very low *RH2* expression has been observed in tarpon and gobies. In all these fishes, green sensitivity has been taken over by a shorter-wavelength shifted copy of the red sensitive *LWS* opsin (also see Cortesi et al., 2021; Escobar-Camacho et al., 2020; Liu et al., 2018). SWS1 and 2, short-wavelength-sensitive 1 and 2 opsin genes; RH2, rhodopsin-like 2 opsin gene; LWS, long-wavelength-sensitive opsin gene.

The phylogenetic reconstruction showed that the teleost ancestor most likely possessed only one copy of *RH2*. Alternatively, *RH2* duplicates that could have existed in these early fishes, might have been lost over evolutionary time and we are no longer able to detect them in extant species. The first ancestral duplication that we find in our dataset dates to the ancestor of Clupeocephala i.e., after the split of the Elopomorpha and Osteoglossomorpha lineages. At this point, at least two but possibly even three rounds of gene duplications gave rise to four ancestral *RH2* copies (Fig. 1). Two of these, *RH2-II* and *RH2-III*, are only found in the Otomorpha lineage (herring, zebrafish, carp, knifefish, catfish and relatives) and were lost in euteleosts (Figs. 1 and 2). A third copy (*RH2-I*) is found in a limited number of species belonging to phylogenetically very distinct lineages including zebrafish (Cypriniformes, Otomorpha), milkfish (Gonorynchiformes, Otomorpha), smelts (Osmeriformes, Euteleostei), and squirrel and soldierfishes (Holocentriformes, Euteleostei). The fourth ancestral copy, by convention referred to as *RH2*, served as the starting point for the immense diversification of the green-sensitive opsins in euteleosts (Figs.1 and 2). Hence, the onset of the *RH2* duplications, or more likely the surviving ancestral duplicates are from a different evolutionary time point compared to the other visual opsins (Musilova et al., 2021). Both, the ancestral duplicates of *RH1* (Chen et al., 2018; Musilova et al., 2019) and *LWS* (Cortesi et al., 2021) can be dated back to the teleost ancestor or even earlier. On the contrary, the surviving ancestral *SWS2* duplicates first occurred later during the teleost evolution, in the neoteleost and percomorph ancestors (Cortesi et al., 2015).

Following the ancestral duplication events, the evolution of *RH2* appears even more dynamic with numerous gene duplications and gene losses that occurred also in more recent evolutionary times (Fig. 1 and Fig. S1). For example, differences between species of the same order such as in the Carangiformes, Gobiiformes, and Clupeiformes are quite common (Fig. S1). Interestingly, while in the Otomorpha this diversity is based on all four ancestral duplicates (Fig. 2A), in euteleosts most of the *RH2* diversity stems from subsequent duplications of the fourth ancestral *RH2* copy (Fig. 1). First, a gene duplication in the Neoteleostei ancestor gave rise to the *RH2A* gene. The second copy from this event then duplicated again in the ancestor of acanthopterygians, giving rise to the *RH2B* and *RH2C* copies (Fig. 1). The occurrence of the *RH2A* and *RH2B* copies is common in percomorph fishes and has been widely reported in previous opsin gene studies (reviewed in Lin et al., 2017; Musilova et al., 2021; Rennison et al., 2012). In this study we identify for the first time a third percomorph-specific copy, *RH2C*, which is present in numerous species including tunas, trevallies, labrids and sunfish (Fig. 1 and Fig. S1). *RH2C* was likely lost in the ancestor of Ovalentaria, and this might explain why it has been overlooked in previous studies as it is not found in many of the commonly studied model species for vision such as medaka (Matsumoto et al., 2020), cichlids (Carleton and Yourick, 2020), killifish (Chang et al., 2021) and dottybacks (Cortesi et al., 2016). Finally, pikes (Esociformes) and salmonids (Salmoniformes), the first lineages to split from the euteleosts, have undergone independent duplications of the fourth ancestral *RH2* copy (Figs. 1 and 2B) and do not share *RH2* paralogs with the otomorphs (*RH2-I, II* and *III*) or the neoteleosts (*RH2A, B* and *C*).

Most of the *RH2* genes are found within the same genomic clusters (data not shown). There are two main *RH2* synteny groups in teleosts. In the majority of euteleost species all *RH2* genes occur in tandem on the same chromosomal cluster flanked by the *synpr* and *slc6A13* genes. The Otomorpha *RH2-I* and *RH2-III* copies are also located in the same cluster with the same flanking genes, while the *RH2-II* copy is located in a different cluster next to *mutS-H5*. In some species which experienced whole genome duplications there are two highly similar *RH2* clusters (e.g., in goldfish, *Carassius auratus*). However, only a single *RH2* cluster is present in Salmoniformes, despite salmon also having undergone a lineage specific whole genome duplication (Ss4R) (Lien et al., 2016). In very few cases such as in darters (*Etheostoma* spp.) and pike (*Esox lucius*), we found that their *RH2* clusters were surrounded by different genes. In these instances, it is likely that genomic rearrangements have affected the *RH2* clusters. Alternatively, *RH2* copies might have moved around the genome with the help of transposable elements before subsequent duplications took place.

*RH2* genes also appear strongly impacted by gene conversion (reviewed in Musilova et al., 2021). Gene conversion, that is the unidirectional exchange of genetic information between similar gene copies, may homogenise opsin gene coding regions by replacing parts of the original gene with its paralog (e.g., Cortesi et al., 2015; Sandkam et al., 2017). Two ancestrally differentiated copies might then appear like recently duplicated genes in the gene trees, as observed in our dataset. For example, the three *RH2s* (two *RH2A* and one *RH2B*) found in many percomorphs might have a common origin, but the synteny is not consistent between species; in some cases one of the genes is arranged in the reverse directions other times, it is not (Musilova et al., 2019) (Fig. 1). To resolve the true relationship between *RH2* paralogs that have experienced gene conversion, several studies took the approach of removing the converted parts of the genes before tree reconstruction. This has shown that for example the *RH2A* paralogs found in Amazonian cichlids have a shared ancestry with the *RH2A* paralogs in African and Neotropical cichlids (Escobar-Camacho et al., 2017). Also, in the spotted unicornfish, *Naso brevirostris*, removing the converted regions recovered the sister relationship between its *RH2A* and *RH2B* copies (Tettamanti et al., 2019). In general, gene conversion appears to be an important evolutionary mechanism for visual opsins [e.g., for *SWS2* (Cortesi et al., 2015), and *LWS* (Cortesi et al., 2021; Escobar-Camacho et al., 2020; Sandkam et al., 2017)], that quite possibly assists in keeping their function restricted to a certain spectral-sensitivity range.

Indeed, using gene expression from retinal transcriptomes in 30 species representing the breath of the teleost phylogeny, we show that *RH2* is the dominantly expressed green sensitive opsin gene in most teleost species (Fig. 3). This highlights the importance of *RH2* opsins for vision in different habitats ranging from clear streams over murky lakes to the relative darkness of the deep sea. However, in species which have lost (Osteoglossomorpha and some of the characins) or do not express (most characins and gobies) *RH2*, shorter wavelength shifted copies of the *LWS* opsin have taken up the green sensitive niche (also see Liu et al. 2018, Escobar-Camacho et al. 2020, and Cortesi et al. 2021).

It is important to note that it is the recent advances in whole genome sequencing and particularly the high-quality genomes produced by long-read sequencing and advanced assembly pipelines (e.g., Rhie et al., 2021), which have made it possible to gain a thorough overview of the evolutionary history of the teleost *RH2* opsin genes. Conventional short-read based assemblies often failed in the regions with multiple *RH2*s due to the high similarity between copies, numerous gene duplicates and frequent gene conversion. By looking at over 100 fish genomes, our study provides a comprehensive first insight into the *RH2* evolution in fishes. However, we expect future work to resolve many more of the evolutionary intricacies that are hidden away in the genomes of the more than 34,000 teleost species.

## Material and Methods

### Mining of RH2 opsins from the whole genome dataset

Whole genomic data have been downloaded from GenBank for the species with high-quality genomes. In a few cases, we included lower quality draft genomes if the *RH2* genes were assembled as one cluster or resembling the synteny in high quality genomes. Table S1 contains the details for all species including accession numbers.

To search for the *RH2* opsin genes, we have used the following pipeline: assembled genomes have been mapped against the single exons 1 and 4 of the eel (*Anguilla japonica*), blind cavefish (*Astyanax mexicanus*), zebrafish (*Danio rerio*; *RH2-I* and *RH2-III*), pike (*Esox lucius*), squirrelfish (*Neoniphon sammara; RH2-I, RH2A* and one uncertain copy) and tilapia (*Oreochromis niloticus*; *RH2Aalpha* and *RH2B*). Mapping was performed in Geneious 9.1.8 (www.geneious.com) using the Medium Sensitivity settings. This setting was sensitive enough to make hits on all *RH2* and *RH1* and exo-rhodopsin, the latter two have subsequently been excluded. Scaffolds/chromosomes with positive hits were then screened more in more detail. First, all single exons of all opsin genes from the species mentioned above have been mapped against the scaffold/chromosome with the High Sensitivity settings. This has forced the exons to be mapped in the regions with opsins, and as such served to identify the *RH2* cluster. Once the cluster(s) have been found, we inspected the upstream and downstream region (by mapping again the single exons) to make sure there are no more *RH2* genes in this scaffold. In the *RH2* cluster, we mapped our exons also to every intergenic region (separately), to make sure no gene fragment was overlooked. After extracting the *RH2* genes, we annotated them in Geneious using the “automated annotation” function and the exon/intron boundaries in the annotation were subsequently confirmed via single exon mapping. In those cases, in which annotated genomes were available we used those annotations, but we nevertheless doublechecked the upstream and intergenic regions to exclude the presence of further *RH2* genes.

### Phylogenetic analysis

The final dataset contained 417 *RH2* sequences (plus one exo-rhodopsin outgroup). All *RH2* sequences were aligned using the MAFFT plug-in as implemented in Geneious v.9.1.8 (Katoh et al., 2019). Due to the uncertainties with the identity and annotation of exon 5 (shorter than other exons), we have excluded this exon from the alignment. We have reconstructed the *RH2* gene tree using the MrBayes 3.2 (Ronquist et al., 2012) software on the CIPRES portal (Miller et al., 2010). MrBayes was run four times independently (each with two runs) with 50 million generations each. We the visually inspected the resulting output files in Tracer 1.5 (https://beast.community/) to identify the run with the best -lnL score and the number of generations it took for each run to reach the -lnL plateau. Four out of the eight resulting files were selected, and a consensus tree reconstructed using a burnin of 25 million generations.

### Transcriptome analysis

Transcriptomic reads were mapped against a general reference including all known rod and cone opsin gene types (references from: tilapia, zebrafish, Round goby, cavefish) with Medium Sensitivity settings in Geneious 9.1.8. If only one copy per type was found in a given species, the consensus was exported and used as a species-specific reference. If multiple copies per opsin type were present, we manually disentangled the copies (for details methods see de Busserolles et al., 2017 and Musilova et al., 2019). The transcriptomic reads were then mapped again against each species-specific reference dataset with the Lowest Sensitivity settings. The number of reads that mapped to each copy were then used to calculate the proportion (in %) of expression of each opsin type from the total cone opsin expression as per de Busserolles et al., 2017 and Tettamanti et al., 2019.

## Acknowledgement

We would also like to acknowledge the indigenous owners of the land on which some of the reported research has been carried out or from which specimens reported in this study derive. FC was supported by an Australian Research Council DECRA Research Fellowship (ARC DE200100620) and a University of Queensland Development Fellowship. ZM was supported by the Swiss National Science Foundation (SNF grant PROMYS, 166550), Charles University (Primus), and the Czech Science Foundation (21-31712S).

---

**Supplementary Figure S1:**
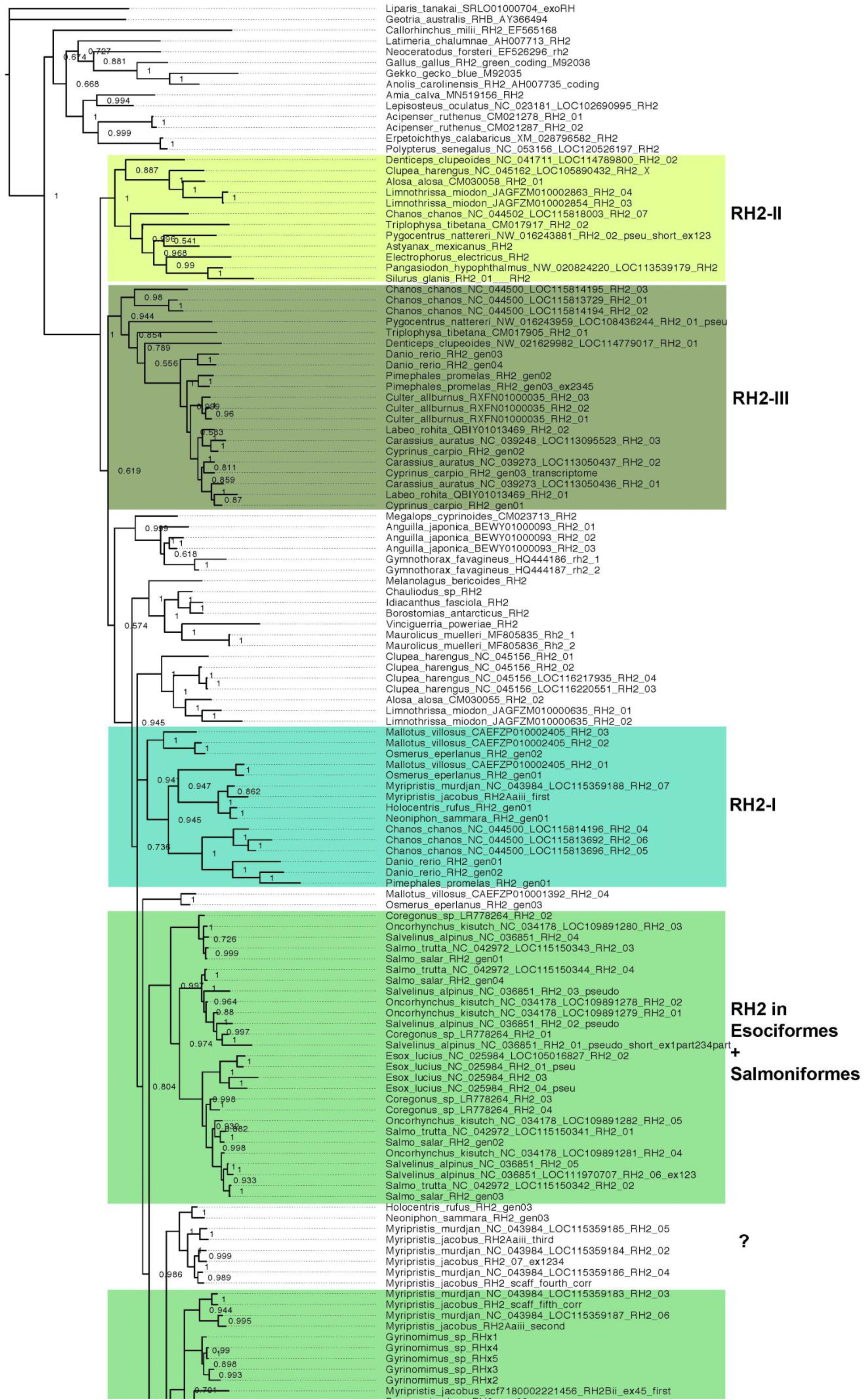
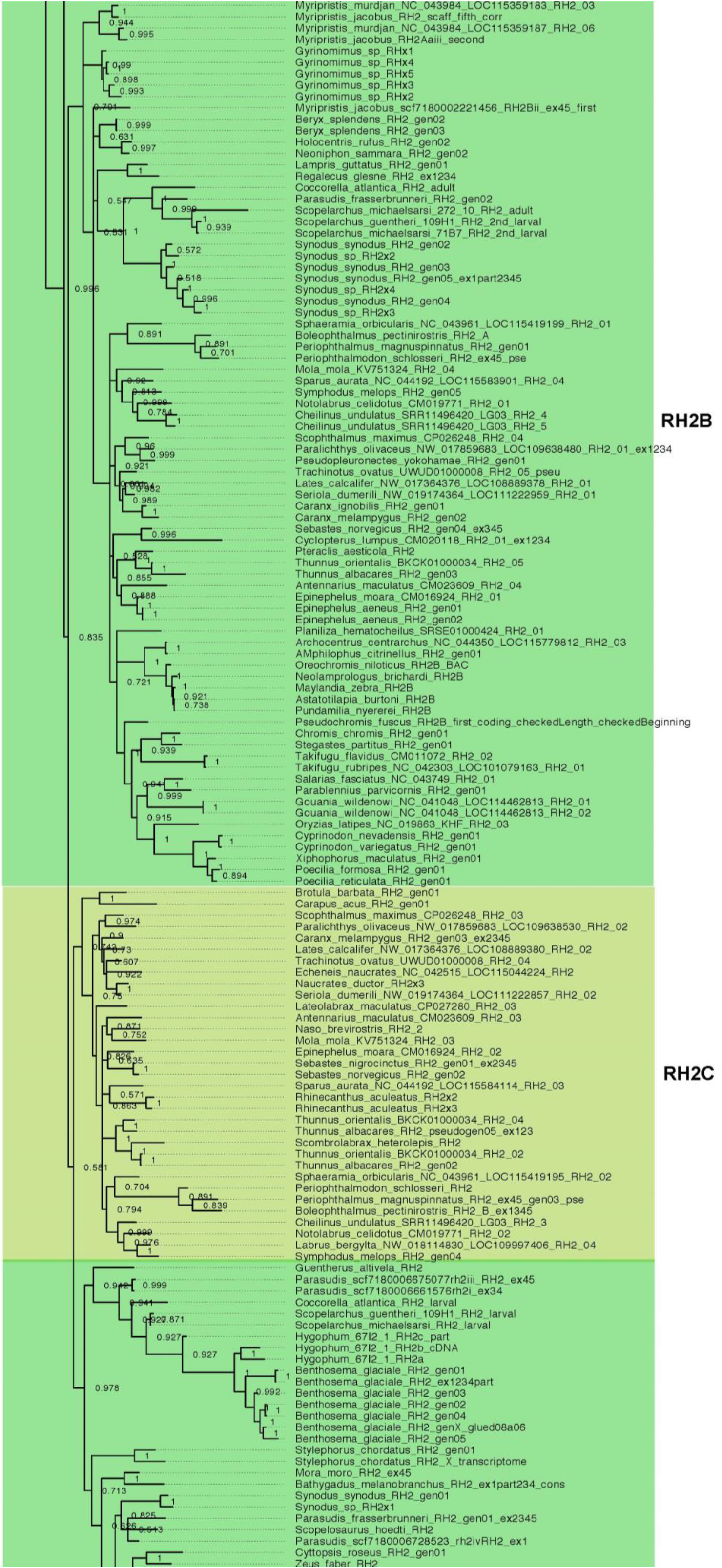
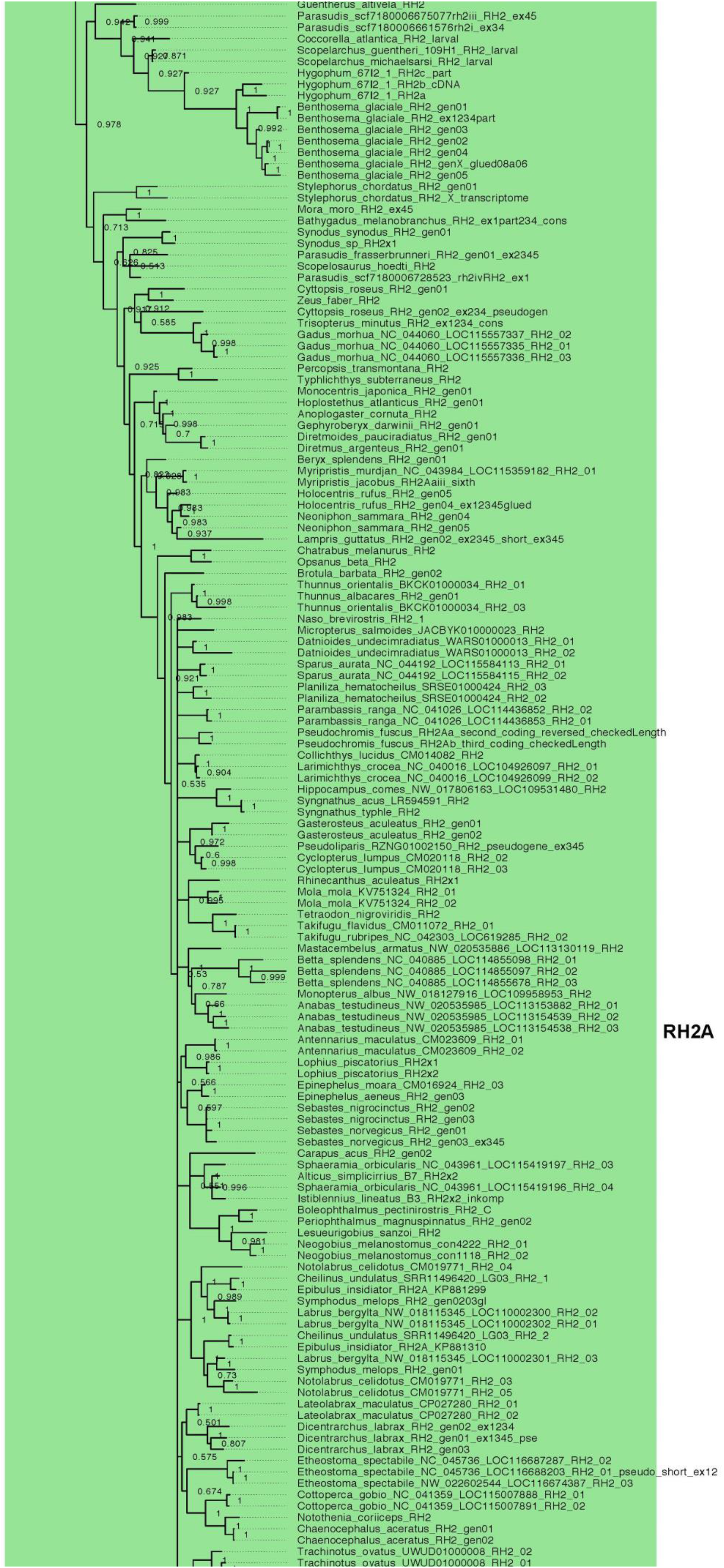
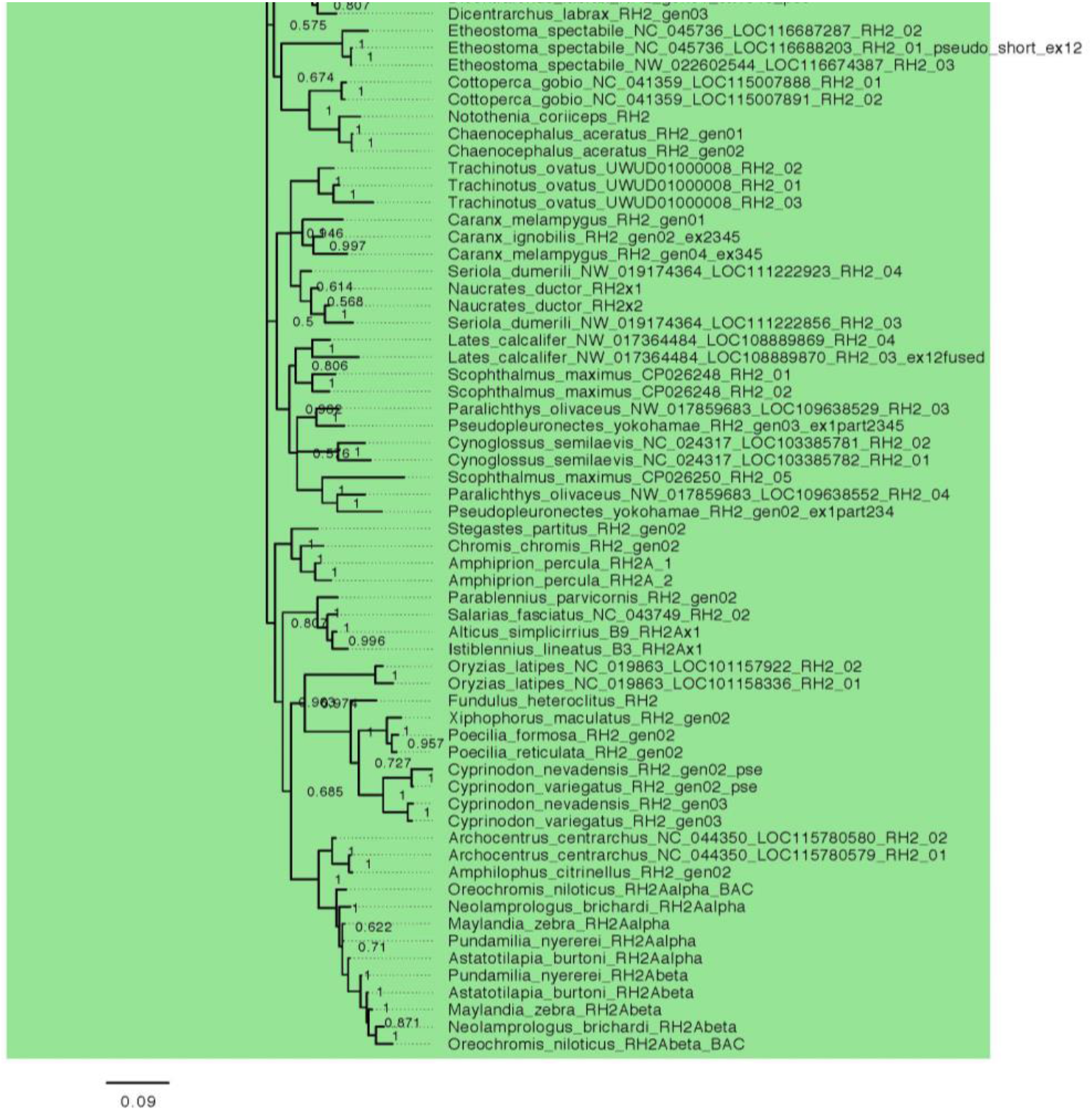
*RH2* gene tree including all sequences mined from the whole genome dataset of 115 teleost species and three non-teleost outgroups.

